# A generic hidden Markov model for multi-parent populations

**DOI:** 10.1101/2021.08.03.454963

**Authors:** Karl W. Broman

## Abstract

A common step in the analysis of multi-parent populations is genotype reconstruction: identifying the founder origin of haplotypes from dense marker data. This process often makes use of a probability model for the pattern of founder alleles along chromosomes, including the relative frequency of founder alleles and the probability of exchanges among them, which depend on a model for meiotic recombination and on the mating design for the population. While the precise experimental design used to generate the population may be used to derive a precise characterization of the model for exchanges among founder alleles, this can be tedious, particularly given the great variety of experimental designs that have been proposed. We describe an approximate model that can be applied for a variety of multi-parent populations. We have implemented the approach in the R/qtl2 software, and we illustrate its use in applications to publicly-available data on Diversity Outbred and Collaborative Cross mice

## Introduction

Multi-parent populations (MPPs) are valuable resources for the analysis of complex traits (de Koning and McIntyre 2017), including the mapping of quantitative trait loci (QTL). A wide variety of MPPs have been developed, including heterogeneous stock (HS) in mice (Mott *et al*. 2000) and rats (Solberg Woods *et al*. 2010), eight-way recombinant inbred lines (RIL) in mice (Complex Trait Consortium 2004) and Drosophila (King *et al*. 2012), and multi-parent advanced generation intercross (MAGIC) populations in a variety of plant species including arabidopsis (Kover *et al*. 2009), wheat (Cavanagh *et al*. 2008), maize (Dell’Acqua *et al*. 2015), and rice (Bandillo *et al*. 2013).

QTL mapping in MPPs can be performed through statistical tests at individual single nucleotide polymorphisms (SNPs), as used in genome-wide association studies. However, many investigators first seek to reconstruct the mosaic of founder haplotypes along the chromosomes of MPP individuals and use this reconstruction to test for association between founder alleles and the quantitative phenotype. This approach was first introduced by Mott *et al*. (2000) for the analysis of HS mice, implemented in the HAPPY software, and has been continued in packages such as R/mpMap (Huang and George 2011), DOQTL (Gatti *et al*. 2014), and R/qtl2 (Broman *et al*. 2019a).

The process of genotype reconstruction in an MPP individual is illustrated in Fig. **1**. The genotypes in the founder strains (Fig. **1a**) and the MPP offspring (Fig. **1b**) are used to calculate the probability of each possible founder genotype at each position along the chromosome (Fig. **1c**). Thresholding of these probabilities can be used to infer the founder genotypes and the locations of recombination breakpoints (Fig. **1d**).

**Figure 1.**
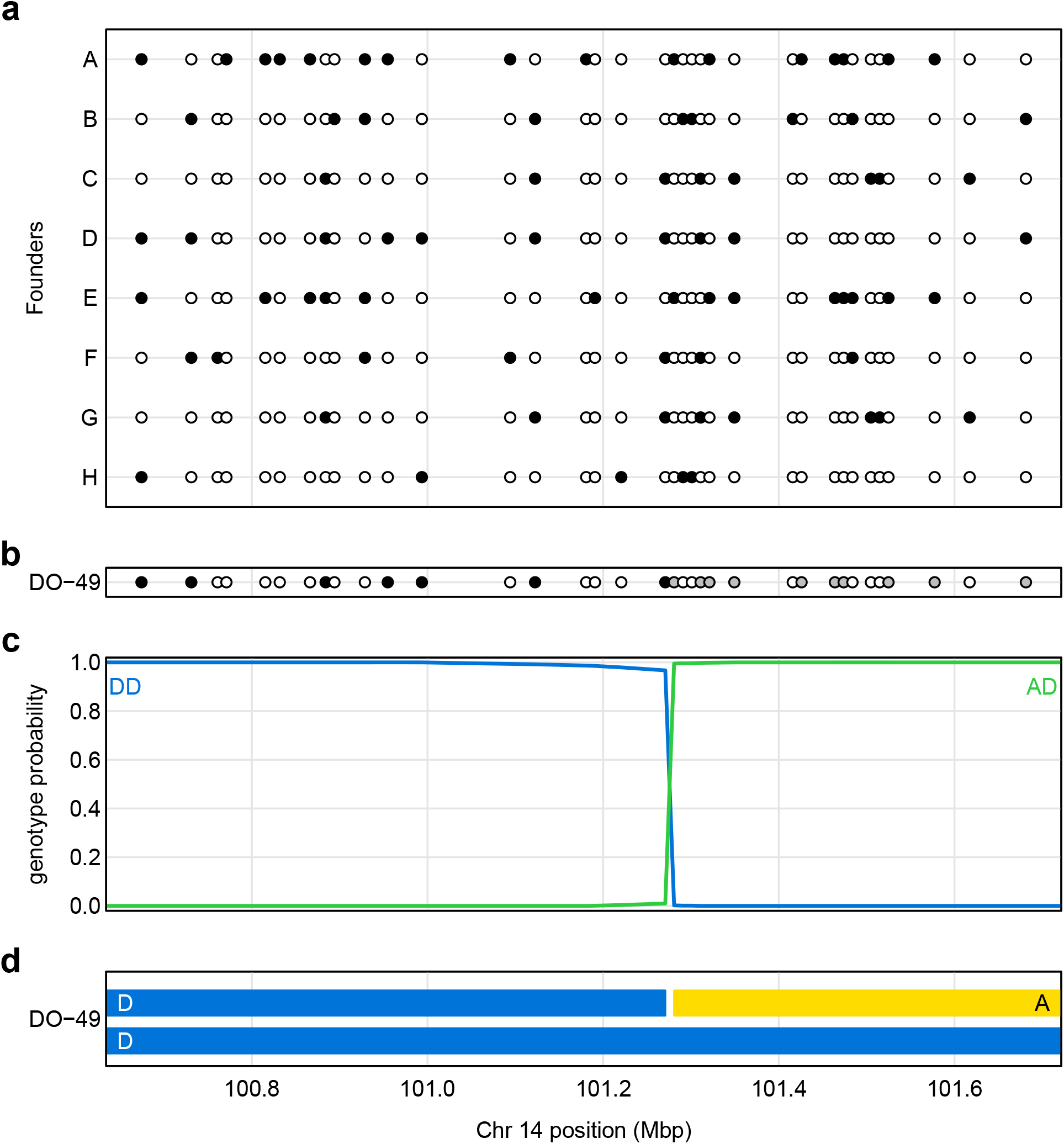
Illustration of genotype reconstruction in a 1 Mbp region in a single Diversity Outbred (DO) mouse. **a**. Genotypes of eight founder strains at a set of SNPs, with open and closed circles corresponding to being homozygous for the more-frequent and less-frequent allele, respectively. **b**. Genotype of the DO mouse at the SNPs, with gray indicating the mouse is heterozygous.**c**. Genotype probabilities for the DO mouse along the chromosome segment, given the observed data. Genotypes other than the two shown have negligible probability across the region. **d**. Inferred haplotypes in the DO mouse.

Such genotype reconstructions are valuable not just for QTL analysis but also for data diagnostics (Broman *et al*. 2019b). For example, the inferred number of recombination breakpoints is a useful diagnostic for sample quality. Further, the reconstructed genotypes can be used to derive predicted SNP genotypes; comparing these to the observed SNP genotypes can help to identify problems in both samples and SNPs.

The probability calculation in Fig. **1c** depends on a model for the process along MPP chromosomes in Fig. **1d**. In the HAPPY software for HS mice, Mott *et al*. (2000) used a model of random mating in a large population. Broman (2005) extended the work of Haldane and Waddington (1931) to derive two-locus genotype probabilities in multi-parent recombinant inbred lines. This was later developed for the case of multi-parent advanced intercross populations (Broman 2012a, b), including Diversity Outbred (DO) mice (Churchill *et al*. 2012).

Genotype reconstruction for a variety of MPP designs has been implemented in the R/qtl2 software (Broman *et al*. 2019a, https://kbroman.org/qtl2). But it can be tedious analytical work to derive the appropriate transition probabilities for each new MPP design that is proposed. An alternative is to develop a more general approach for genotype reconstruction, such as used in the software RABBIT (Zheng *et al*. 2015). However, this approach has a variety of parameters that can be difficult to specify.

Here we propose a similarly general method for genotype reconstruction in MPPs. We imagine that an MPP was derived from a population of homozygous founder strains at known proportions, *α_i_*, followed by *n* generations of random mating among a large number of mating pairs. We can derive the exact transition probabilities for this situation. The *α_i_* should be simple to specify from the MPP design, and the effective number of generations of random mating, *n*, can be determined by computer simulation, to match the expected density of recombination breakpoints.

Our approach has been implemented in R/qtl2. While we currently focus on data with SNP genotype calls, such as from microarrays, our model could potentially be incorporated into methods for genotype imputation from low-coverage sequencing, such as that of Zheng *et al*. (2018). We illustrate our approach through application to publicly-available datasets on DO (Al-Barghouthi *et al*. 2021) and Collaborative Cross mice (Srivastava *et al*. 2017).

## Methods

For genotype reconstruction in a multi-parent population (MPP), we use a hidden Markov model (HMM; see Rabiner 1989). Our basic approach is as described in Broman and Sen (2009, App. D) for a biparental cross; the extension to an MPP is straightforward and described below.

Consider an MPP derived from *k* inbred lines. We focus on a single individual, and on a single chromosome with *M* marker positions (including pseudomarkers: positions between markers at which we have no data but would like to infer the underlying genotype). Let *G_m_* be the underlying genotype at position *m*. In a homozygous population, such as RIL, the *G_m_* take one of *k* possible values, the *k* homozygous genotypes. In a heterozygous population, such as advanced intercross lines (AIL), the *G_m_* take one of 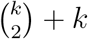 possible values, the 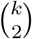 heterozygotes and *k* homozygotes. Let *O_m_* be the observed SNP genotype at position *m* (possibly missing). We assume that the *G_m_* form a Markov chain (that *G*_1_,…, *G*_*m*−1_ are conditionally independent of *G*_*m*+1_,…, *G_m_*, given *G_m_*), and that *O_m_* is conditionally independent of everything else, given *G_m_*. The forward-backward algorithm (see Rabiner 1989) takes advantage of the conditional independence structure of the HMM to calculate Pr(*G_m_*|**O**).

The key parameters in the model are the *initial* probabilities, *π_g_* = Pr(*G*_1_ = *g*), the *transition* probabilities, *t_m_* (*g, g*′) = Pr(*G*_*m*+1_ = *g* | *G_m_* = *g*), and the *emission* probabilities, *e_m_*(*g*) = Pr(*O_m_*|*G_m_* = *g*). A particular advantage of the HMM for genotype reconstruction is the easy incorporation of a model for genotyping errors (Lincoln and Lander 1992), which is done through the emission probabilities, which condition on the founder SNP genotypes but allow some fixed probability ∊ that the observed SNP genotype in the MPP individual is in error and incompatible with the underlying genotype *G_m_* and the SNP genotypes in the founder lines.

The initial and transition probabilities govern the underlying Markov chain, including the relative frequency of founder alleles and the frequency of recombination breakpoints along MPP chromosomes. In principle, these probabilities may be derived on the basis of the crossing design for the MPP. In practice, the transition probabilities can be tedious to derive, and exact calculations may provide no real advantage for genotype reconstruction.

Here, we derive the transition probabilities for a generic MPP design, which may then be applied generally. We consider a founder population with *k* inbred lines in proportions *α_i_*, and imagine subsequent generations are produced by random mating with a very large set of mating pairs.

Consider a pair of loci separated by a recombination fraction of *r* (assumed the same in both sexes) and let 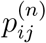 be the probability of that a random haplotype at generation *n* has alleles *i* and *j*. At *n* = 0, we have just the founding inbred lines, and so 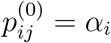 if *i* = *j* and = 0 if *i* ≠ *j*.

The probabilities from one generation to the next are related by a simple recursion, as in Broman (2012b). Consider a random haplotype at generation n. It was either a random haplotype from generation *n* – 1 transmitted intact without recombination, or it is a recombinant haplotype bringing together two random alleles. Thus

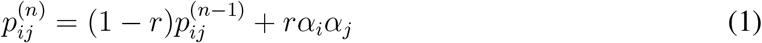

Using the same techniques described in Broman (2012b), we find the solutions:

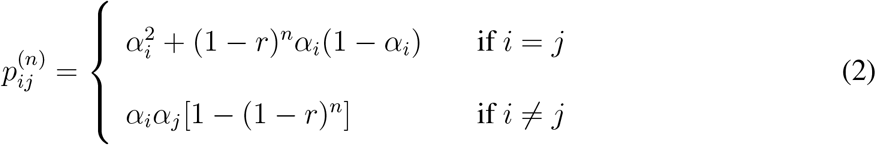

The transition probabilities along a haplotype are derived by dividing the above by the marginal probability, *α_i_*. Thus if *G*_1_ and *G*_2_ are the genotypes at the two loci, we have the following transition probabilities.

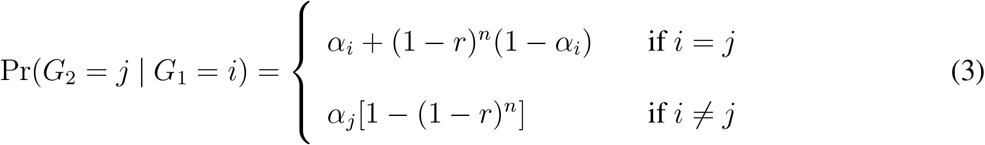

For a heterozygous population (such as heterogeneous stock or Diversity Outbred mice), an individual will have two random such haplotypes. For homozygous population (such as MAGIC), we treat them like doubled haploids, by taking a single random chromosome and doubling it.

For the X chromosome, we use the same equations but replace *n* with (2/3)*n*, since recombination occurs only in females, so in 2/3 of the X chromosomes. This provides a remarkably tight approximation.

You can potentially use the expected number of crossovers to calibrate the number generations of random mating, or the map expansion, which is the relative increase in the number of crossovers. Let *R*(*r*) be the chance that a random haplotype has an exchange of alleles across an interval with recombination fraction *r*, so 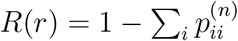. The map expansion is *dR/dr* evaluated at *r* = 0 (see Teuscher and Broman 2007). Using equation (2) above, we then get that the map expansion in this population is 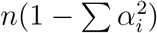. In the special case that *α_i_* ≡ 1/*k* for all *i*, this reduces to *n*(*k* – 1)/*k*.

The map expansion at generation *s* in DO mice on an autosome is (7/8)(*s* – 1) + *M*_1_ where *M*_1_ is the weighted average of map expansion in the pre-CC founders (Broman 2012b), or about (7*s* + 37)/8. Equating this with (7/8)*n*, we can thus take *n* ≈ *s* + 5 when using this model to approximate the DO. For the Collaborative Cross, Broman (2005) showed that R = 7r/(1+6r), and so the map expansion is 7. Thus we can take n = 8 as the effective number of generations of random mating.

## Applications

We illustrate our approach with application to datasets on Diversity Outbred mice (Al-Barghouthi *et al*. 2021) and Collaborative Cross (CC) mice (Srivastava *et al*. 2017). In both cases, the approach provided results that were generally equivalent to those from the more exact model, though with important differences in the results for the X chromosome in the CC application.

### Diversity Outbred mice

The Diversity Outbred mouse data of Al-Barghouthi *et al*. (2021) concerns a set of 619 mice from DO generations 23–33, in 11 batches by generation and including 304 females and 315 males. The mice were genotyped on the GigaMUGA array (Morgan *et al*. 2016) and the cleaned data consist of genotypes at 109,427 markers. A wide variety of phenotypes are available; we focus on the 20 contributing to the results in Table 1 of Al-Barghouthi *et al*. (2021).

We performed genotype reconstruction using the transition matrices derived specifically for DO mice (Broman *et al*. 2019b) as well as by the approximate model proposed above. For the DO mice at generation n, we used the transition probabilities for general 8-way advanced intercross lines (AIL) at n + 5.

Following Al-Barghouthi *et al*. (2021), we assumed a 0.2% genotyping error rate and used the Carter-Falconer map function (Carter and Falconer 1951). Calculations were performed in R (R Core Team 2021) with R/qtl2 (Broman *et al*. 2019a), on an 8-core Linux laptop with 64 GB RAM. The calculations with the DO-specific model took approximately 35 min, while those with the general AIL model took 27 min, an almost 25% reduction in computation time.

The transition probabilities used by the two models are only subtly different and become less different in later generations. The probability of an exchange across an interval on a random DO chromosome, as a function of the recombination fraction for the interval and the number of generations, is shown in Fig. **2**.

**Figure 2.**
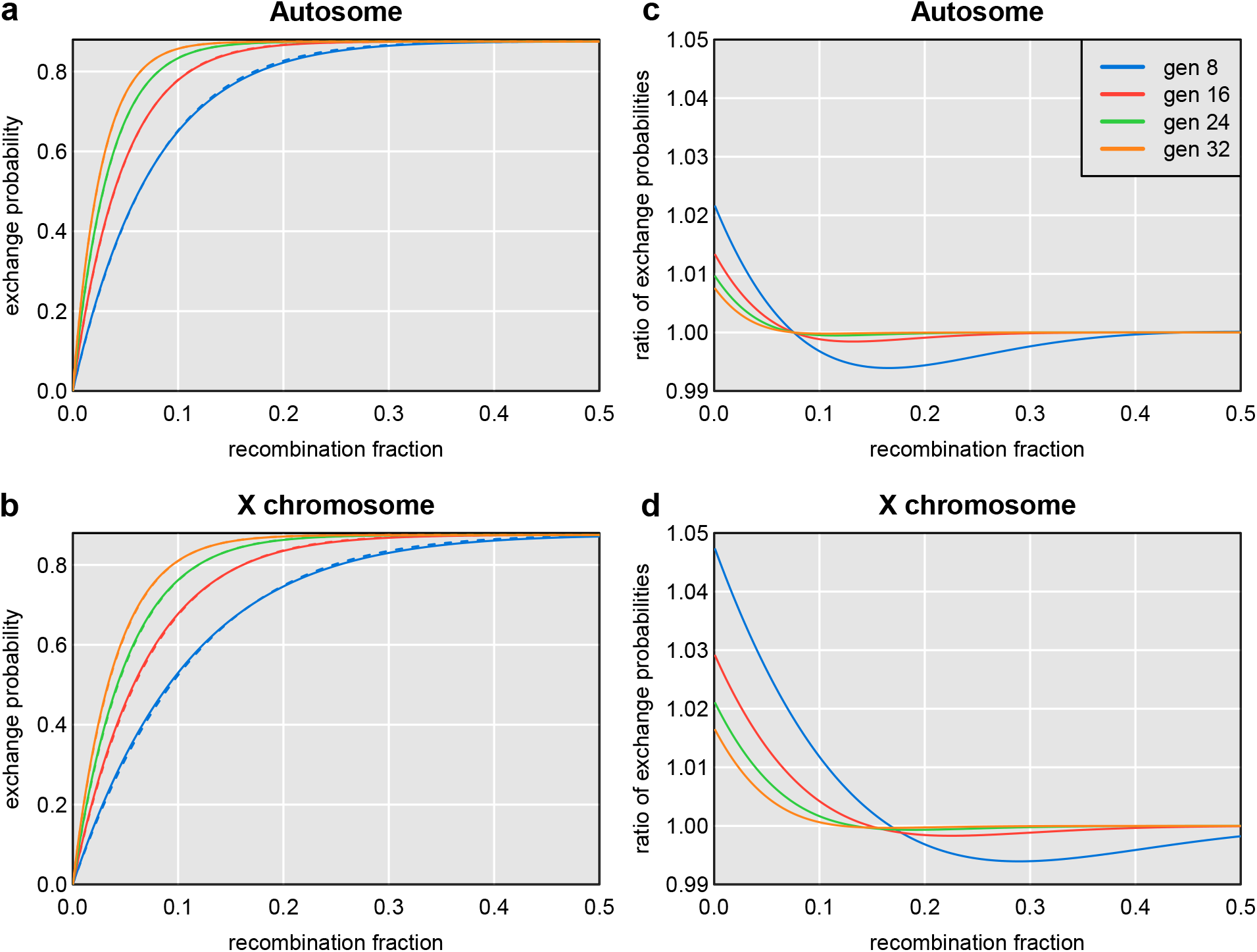
Differences in transition probabilities for Diversity Outbred mice from more-exact calculations and the proposed approximations. Probability of an exchange of alleles across an interval as a function of generation with the more-exact calculations (solid lines) and the proposed approximation (dashed lines) for autosomes (**a**) and the X chromosome (**b**). Ratio of the probabilities (more-exact versus approximation) for autosomes (**c**) and the X chromosome (**d**).

QTL analysis proceeded by the method described in Gatti *et al*. (2014) and also used by Al-Barghouthi *et al*. (2021). Namely, we fit a linear mixed model assuming an additive model for the founder haplotypes, with a residual polygenic effect to account for relationships among individuals with kinship matrices calculated using the “leave-one-chromosome-out” (LOCO) method (see Yang *et al*. 2014), and with a set of fixed-effect covariates defined in Al-Barghouthi *et al*. (2021).

The genotype probabilities were almost indistinguishable. The maximum difference was 0.011 on the X chromosome followed by a difference of 0.007 on chromosome 8. For that reason, the QTL mapping results were hardly different. Across all 20 traits considered, the maximum difference in LOD scores in the two sets of results was 0.02.

The LOD curves by the two methods for tissue mineral density (TMD) and the differences between them are shown in Fig. **3**. The QTL on chromosomes 1 and 10 have LOD scores of 23.9 and 14.6, respectively, but the maximum difference in LOD, genome-wide, between the two methods is just 0.014.

**Figure 3.**
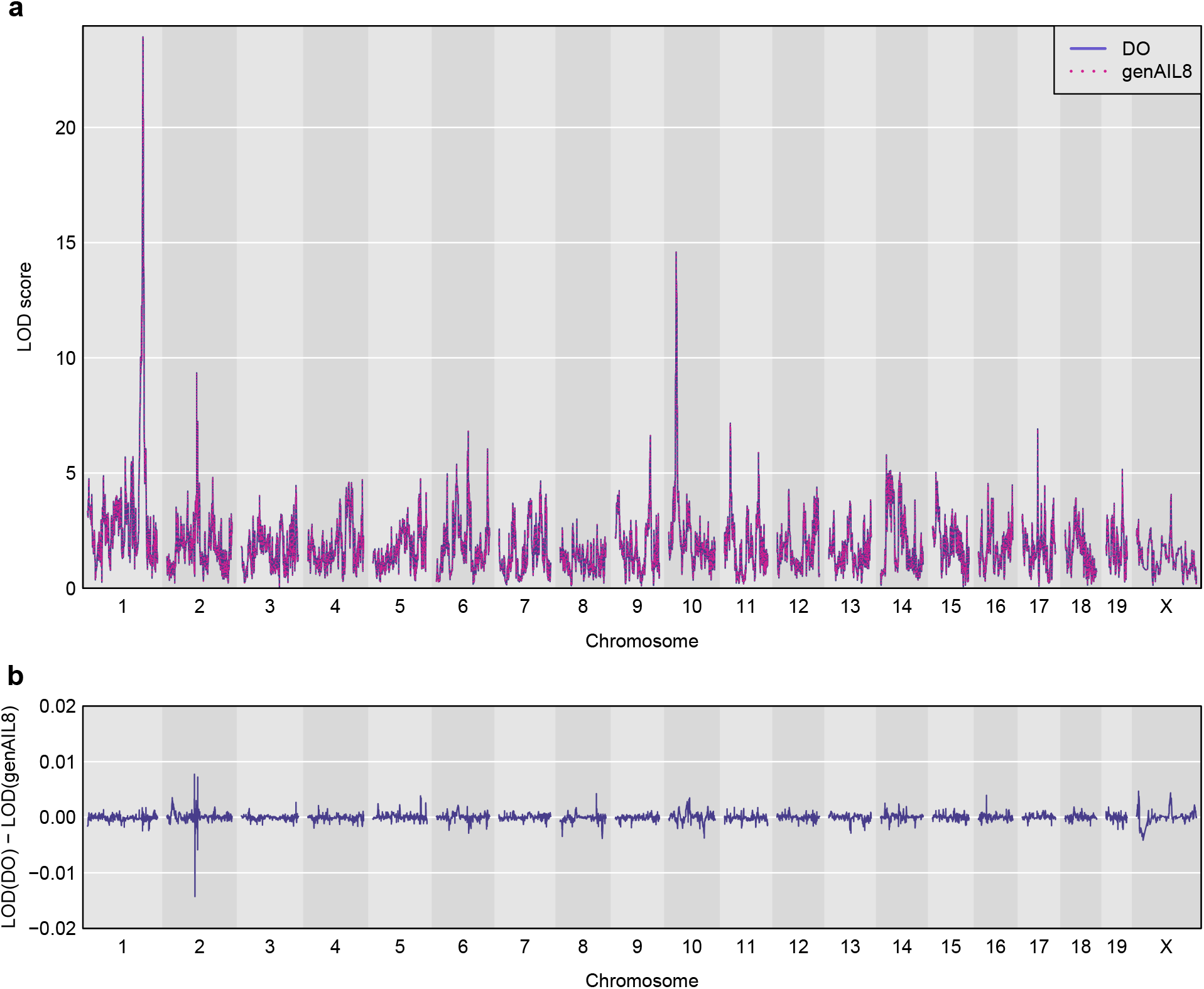
Genome scan for tissue mineral density (TMD) for the DO mouse data from Al-Barghouthi *et al*. (2021). **a**. LOD curves across the genome using the genotype probabilities from the DO-specific model (solid blue curves) and the proposed general model (dotted pink curves). **b**. Differences between the two sets of LOD curves.

### Collaborative Cross mice

As a second application of our approach, we consider the data for a set of 69 Collaborative Cross (CC) lines (Srivastava *et al*. 2017). These are eight-way recombinant inbred lines (RIL) derived from the same eight founders as the DO mice, as the DO was formed from 144 partially-inbred lines from the process of developing the CC (Svenson *et al*. 2012).

Each CC line was formed from a separate “funnel,” bringing the eight founder genomes together as rapidly as possible, for example [(A×B)×(C×D)]×[(E×F)×(G×H)], where the female parent is listed first in each cross. Inbreeding was accomplished by repeated mating between siblings.

The recombination probabilities for the autosomes in the CC do not depend on the order of the founders in the funnel for a line (Broman 2005). This is in contrast with the case of 8-way RIL by selfing (see Broman 2005, Table 2). For the X chromosome, however, the cross order is important, as only 5 of the 8 founders can contribute. For example, in a line derived from the cross [(A×B)×(C×D)]×[(E×F)×(G×H)], the single-locus genotype probabilities on the X chromosome are 1/6 each for alleles A, B, E, and F, and 1/3 for allele C, while alleles D, G, and H will be absent. And note that the mitochondrial DNA will come from founder A, while the Y chromosome will be from founder H.

The cross funnel information was missing for 14 of the 69 CC lines. While the sources of the mitochondria and Y chromosome were provided for all lines, there were several inconsistencies in these data: line CC013/GeniUnc has the same founder listed as the source for its mitochondria and Y chromosome, and for three lines (CC031/GeniUnc, CC037/TauUnc, and CC056/GeniUnc) the founder on the Y chromosome is also seen contributing to the X chromosome. We used the genotype probabilities reported in Srivastava *et al*. (2017) to construct compatible cross funnels, with small modifications to handle the inconsistent information.

We performed genotype reconstruction using the transition matrices derived specifically for CC mice (Broman 2005) as well as by the approximate model proposed above, using *n* = 8 generations of random mating, chosen to match the expected frequency of recombination breakpoints.

The resulting probabilities were nearly identical on all autosomes in all CC lines. The maximum difference in probabilities on the autosomes was just 0.0006.

There were some important differences on the X chromosome, however. There were no cases with high probability pointing to different founder alleles by the two models, but there were several cases where two or more founders cannot be distinguished, but some would be excluded by the assumed cross design.

For example, in Figure **4**, we show the genotype probabilities along the X chromosome for strain CC038/GeniUnc, as calculated with the more-exact model (Figure **4a**) and with the approximate model (Figure **4b**). We also include the results for the case that the more-exact model but when an incorrect cross design was used (Figure **4c**). Note the segment near 135 Mbp, which is inferred to be from founder NOD with the more-exact model but is equally likely B6 or NOD with the approximate model; the B6 and NOD founder strains are identical in the region, but the assumed cross design for the CC038/GeniUnc strain excluded B6. For the results using the incorrect cross design (which excluded not just B6 but also 129 and NOD), the results across the entire chromosome become a chopped-up mess, with an apparent 39 recombination breakpoints, versus 5 when the correct cross information is used.

**Figure 4.**
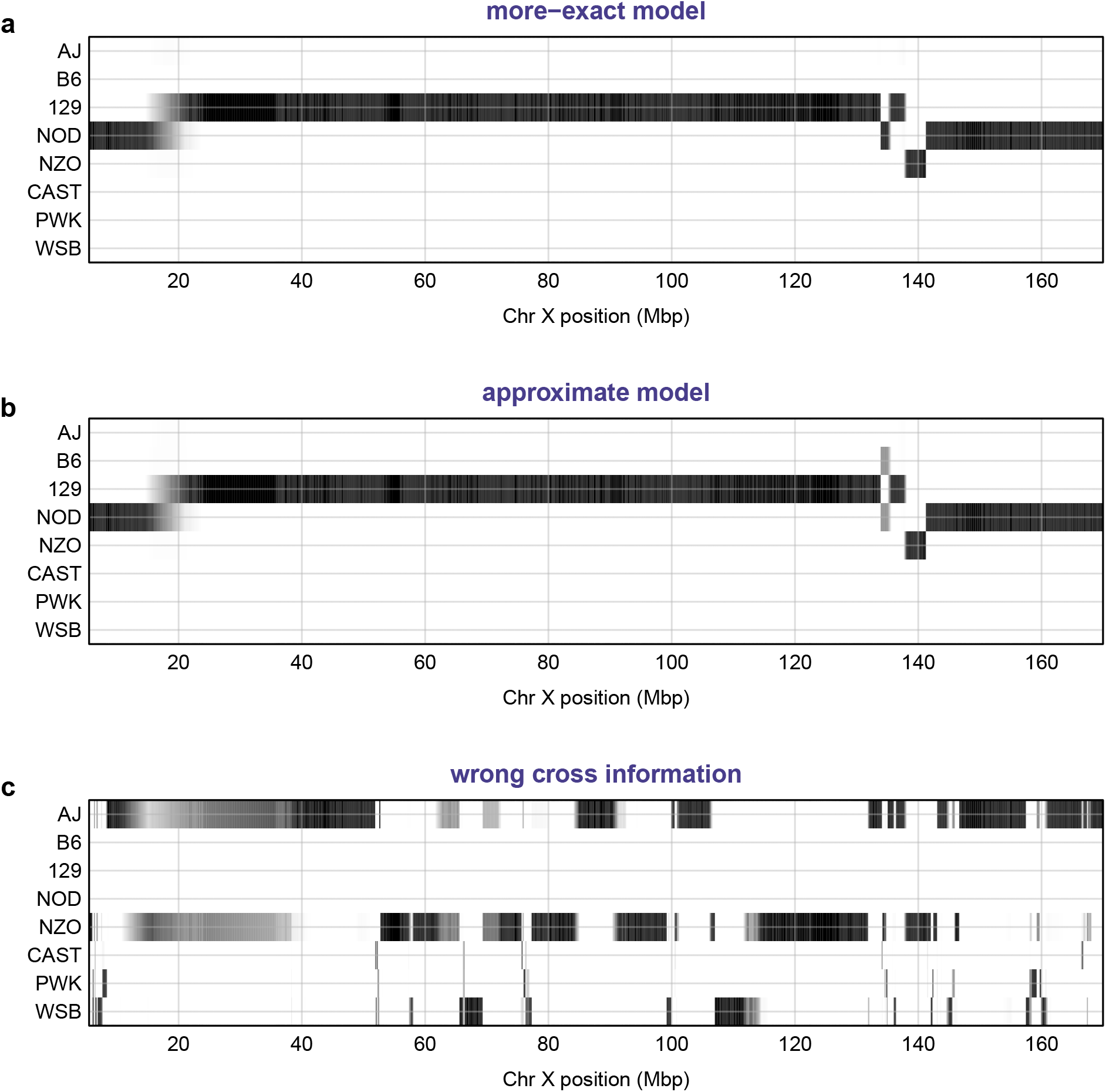
Genotype probabilities along the X chromosome for Collaborative Cross strain CC038/GeniUnc. **a**. Results using the more-exact model that excludes founders B6, CAST, and WSB. **b**. Results using the proposed approximate model. **c**. Results using the more-exact model but with the wrong cross information, excluding founders B6, 129, and NOD.

Overall, there were seven strains where the maximum difference in the probabilities from the more-exact model and the proposed approximate model were in the range 0.25 − 0.50, and another eight strains with maximum difference in the range 0.10 − 0.25. All of the differences concern cases where multiple founders are identical for a region and either some would be excluded by the cross design, or where the difference in prior frequencies affects the results. For example, in the cross [(A×B)×(C×D)]×[(E×F)×(G×H)], the frequency of the C allele on the X chromosome is twice that of A, B, E, and F.

## Discussion

We have proposed an approximate model for use with genotype reconstruction in multi-parent populations (MPPs). We derived the two-point probabilities on autosomes in the case of random mating in large, discrete generations, derived from a founder population of a set of inbred lines in known proportions. We use the same frequencies for the X chromosome, but with 2/3 the number of generations. The approach is shown to give equivalent results for the mouse DO and CC populations, though with important differences for the X chromosome in CC lines, where some founder alleles can be excluded based on the cross design. The more-exact model for the X chromosome in the CC excludes three of the eight founders based on the cross design. This is particularly useful in cases that multiple founders are identical by descent across a region. However, the approximate model is not affected by errors in the specified cross design (see Figure 4).

The value of this generic model points towards the general usefulness of the original software for multi-parent populations, HAPPY (Mott *et al*. 2000), developed for the analysis of mouse heterogeneous stock. The results may depend on marker density and informativeness, but with a dense set of informative markers, a generic approach can provide good-quality genome reconstructions.

The hidden Markov model itself is an approximation. Meiosis generally exhibits positive crossover interference, but the Markov property is closer to being correct in multi-parent populations with multiple generations of mating, because nearby recombination events come from independent generations. This was apparent in the 3-point probabilities derived by Haldane and Waddington (1931) for two-way RIL and was further explored in Broman (2005) for multi-way RIL.

The proposed method has been implemented in the R/qtl2 software (Broman *et al*. 2019a). It requires specification of the founder proportions and one other parameter (the number of generations of random mating) which governs the frequency of recombination breakpoints. The founder proportions should be straightforward from the cross design; the effective number of generations of random mating may require some calibration, such as through computer simulation to match the expected frequency of recombination breakpoints.

## Data and software availability

The R/qtl2 software is available at the Comprehensive R Archive Network (CRAN), https://cran.r-project.org/package=qtl2, as well as GitHub, https://github.com/rqtl/qtl2. Further documentation is available at the R/qtl2 website, https://kbroman.org/qtl2.

The Diversity Outbred mouse data from Al-Barghouthi *et al*. (2021) is available at Zenodo, https://doi.org/10.5281/zenodo.4265417. Also see their companion repository of analysis scripts at GitHub, https://github.com/basel-maher/DO_project, and archived at Zenodo, https://doi.org/10.5281/zenodo.4718146.

The Collaborative Cross mouse data from Srivastava *et al*. (2017) is available at Zenodo, https://doi.org/10.5281/zenodo.377036. Reorganized files in R/qtl2 format are at https://github.com/rqtl/qtl2data/tree/master/CC.

Our detailed analysis code is available at GitHub, https://github.com/kbroman/Paper_GenericHMM, and archived at Zenodo, https://doi.org/10.5281/zenodo.5156660.

## Acknowledgments

Two anonymous reviewers provided valuable comments for improvement of the manuscript.

## Funding

This work was supported in part by National Institutes of Health grant R01GM070683.

## Conflicts of interest

The author declares that there is no conflict of interest.

## Notes

https://github.com/kbroman/Paper_GenericHMM

### Competing Interest Statement

The authors have declared no competing interest.

### Summary of Updates

- small changes for clarification - weakened the language for some conclusions

